# Circadian Clock Coordination of the Notch-to-Wnt Signaling Switch Promotes Regenerative Myogenesis

**DOI:** 10.64898/2026.10.31.763285

**Authors:** Tali Kiperman, Xuekai Xiong, Jemima Pangemanan, David Horne, Vijay K. Yechoor, Ke Ma

**Affiliations:** Department of Diabetes Complications & Metabolism, Arthur Riggs Diabetes & Metabolism Research Institute, Beckman Research Institute of City of Hope, Duarte, CA 91010; Department of Molecular Medicine, Beckman Research Institute of City of Hope, Duarte, CA 91010; Division of Endocrinology and Metabolism, Diabetes and Beta Cell Biology Center, Department of Medicine, University of Pittsburgh, Pittsburgh, PA, 15213, USA

**Keywords:** muscle stem cells, myogenesis, circadian clock, muscular dystrophy, Notch signaling

## Abstract

Regenerative deficits of muscle stem cells (MuSCs) contribute to progressive muscle loss in muscular dystrophy. Successful regenerative repair requires a Notch to Wnt switch to allow MuSC expansion and myogenic differentiation, yet the underlying mechanisms remain poorly understood. Leveraging complementary genetic and pharmacological approaches, here we identify the circadian clock as a key regulator of the Notch to Wnt signaling transition to promote regeneration with therapeutic potential for dystrophic disease. Circadian clock activity marks cycling myogenic progenitors during muscle regeneration, and clock induction occurs in concert with Notch activation. We identify that key components of the Notch pathway are direct transcriptional targets of the clock that were induced by clock activation. Mechanistically, clock coordination of Notch and Wnt signaling promoted MuSC proliferative expansion while facilitating myogenic progression during regeneration. Importantly, this clock-regulated mechanism was preserved in dystrophin-deficient mouse and human myoblasts, and clock-activating compounds enhanced regenerative myogenesis in dystrophic muscle. Collectively, our findings uncover a circadian clock-controlled mechanism that orchestrates regenerative signaling transition with potential for therapeutic targeting to restore regenerative capacity in dystrophic disease.

## Introduction

Progressive muscle degeneration and impaired regenerative capacity are central features of muscular dystrophy and contribute to the loss of muscle mass and function (Duan, Goemans et al., 2021, Dumont, Wang et al., 2015, Esper, Brun et al., 2025). Muscle stem cells (MuSCs), also known as satellite cells, provide an endogenous source for muscle repair. Their regenerative capacity becomes compromised in dystrophic conditions, limiting effective tissue restoration (Kodippili & Rudnicki, 2023). MuSCs dysfunction also arises from intrinsic defects in stem cell maintenance and myogenic progression, and therefore, is a key driver of regenerative failure in muscular dystrophy (Chang, Chevalier et al., 2016). Successful muscle regeneration requires a coordinated sequence of MuSC activation, proliferative expansion, lineage commitment, and myogenic differentiation to restore damaged muscle fibers (Kuang, Kuroda et al., 2007, Rudnicki, Le Grand et al., 2008). Understanding the molecular mechanisms that coordinate distinct stages of regenerative myogenesis is therefore essential for identifying therapeutic strategies to enhance endogenous muscle repair and preserve muscle function in degenerative muscle diseases.

The circadian clock, an evolutionarily conserved time-keeping mechanism, exerts daily oscillatory control of key biological processes involved in stem cell properties to maintain tissue homeostasis (Benitah & Welz, 2020, Brown, 2014, Janich, Pascual et al., 2011). In skeletal muscle, the core molecular clock, driven by a transcriptional–translational feedback loop, regulates diverse biological functions, including muscle mass maintenance, structural integrity and contractile performance (Andrews, Zhang et al., 2010, Chatterjee & Ma, 2016, Chatterjee, Nam et al., 2013, Chatterjee, Yin et al., 2019, Harfmann, Schroder et al., 2016, Yin, Li et al., 2020). The core clock machinery exerts its pervasive effect on skeletal muscle biology via rhythmic output in gene regulation (Takahashi, 2017). Circadian rhythmic processes in skeletal muscle are enriched in cell cycle regulation, proliferation, and differentiation, suggesting a role in coordinating tissue remodeling (Miller, McDearmon et al., 2007). Beyond maintaining physiological homeostasis, clock-controlled mechanisms regulate stem cell responses to environmental and tissue-derived cues (Janich et al., 2011). In MuSCs, several components of the molecular clock influence proliferative and myogenic properties essential for regenerative repair (Chatterjee et al., 2013, Chatterjee et al., 2019, Chatterjee, Yin et al., 2014, Katoku-Kikyo, Paatela et al., 2021, Lowe, Lage et al., 2018). The essential core clock transcription activator Brain and Muscle Arnt-like 1 (BMAL1) is required for coordinating MuSC proliferative expansion and myogenic differentiation that facilitates regenerative repair (Chatterjee et al., 2013, Chatterjee et al., 2019, Chatterjee, Yin et al., 2015), whereas the clock repressor REV-ERBα negatively regulates regenerative remodeling exerts a negative role that limits regenerative remodeling (Chatterjee et al., 2019, Welch, Billon et al., 2017, Xiong, Gao et al., 2021). Components that constitute the negative feedback arm of the core clock loop, including PER1/PER2 and CRY1/CRY2, also influence myogenesis (Katoku-Kikyo et al., 2021, Lowe et al., 2018). Despite advances in understanding circadian clock function involved in MuSC myogenic behavior, the underlying mechanisms mediating clock activity in distinct stages of myogenic progression remain poorly defined. In particular, how clock-dependent transcriptional programs may orchestrate regenerative events during muscle repair has yet to be elucidated.

In response to muscle injury, an orderly progression of activation, lineage commitment and differentiation of myogenic precursors is essential for regenerative repair to replenish tissue mass (Charge & Rudnicki, 2004). A critical determinant of successful muscle regeneration is the temporal coordination of signaling pathways that govern MuSC proliferation and differentiation. Notch signaling supports satellite cell maintenance and proliferative expansion during early regeneration, whereas subsequent activation of canonical Wnt signaling promotes myogenic differentiation and tissue repair (Brack, Conboy et al., 2008, Brack, Conboy et al., 2007). The transition from Notch to Wnt signaling must be precisely coordinated, as disruption of its timing compromises regenerative progression and muscle repair (Brack et al., 2008, Brack et al., 2007). Previous studies have demonstrated direct circadian transcriptional regulation of canonical Wnt signaling, providing a mechanism for clock-mediated promotion of myogenic differentiation (Chatterjee et al., 2013, Chatterjee et al., 2014). However, whether the circadian clock also regulates Notch signaling or coordinates a Notch-to-Wnt transition remains unknown. Given its intrinsic timing function and transcriptional control of myogenic pathways, the circadian clock represents a potential regulatory mechanism for orchestrating the sequential signaling events required for effective muscle regeneration.

Defective MuSC maintenance and myogenic progression contribute to regenerative dysfunction in muscular dystrophy, where repeated cycles of muscle degeneration and incomplete repair progressively compromise tissue integrity (Duan et al., 2021, Dumont et al., 2015, Esper et al., 2025). Impaired asymmetric division disrupts the balance between self-renewal and differentiation, leading to stem cell depletion, while defective lineage progression compromises the generation of functional myogenic progenitors and mature myofibers (Chang et al., 2016, Esper et al., 2025, Kodippili & Rudnicki, 2023). Elucidating the molecular mechanisms in controlling the regenerative capacity of muscle could lead to targeted interventions to restore the regenerative defects to mitigate the loss of muscle mass and function in dystrophic conditions. Further, therapeutic approaches that restore the coordinated regenerative responses of MuSCs may complement current strategies for muscular dystrophy through correcting the underlying genetic defect or limiting muscle damage. Despite current understandings of clock function involved in muscle regeneration, most studies have relied on genetic loss-of-function models and the therapeutic potential of enhancing clock activity remains largely unexplored. In particular, whether clock activation can restore coordinated regenerative signaling and improve myogenic repair under dystrophic conditions remains unclear. Addressing this gap requires mechanistic investigation using genetic gain-of-function approaches together with pharmacological strategies, with validation in disease-relevant experimental models and human myogenic cells.

In the current study, we combined a novel genetic circadian clock gain-of-function model with small-molecule clock activators to define how clock-dependent transcriptional programs coordinate regenerative myogenesis. We identify the circadian clock as a regulator of the Notch-to-Wnt transition, linking early progenitor expansion to subsequent myogenic differentiation through coordination of regenerative signaling pathways. Importantly, we demonstrate that these clock-regulated mechanisms are preserved in dystrophin-deficient mouse and human myoblasts and that clock-activating compounds promote regenerative myogenesis following acute injury and in dystrophic muscle. Together, our findings uncover a mechanistic link between circadian transcriptional control and muscle regenerative signaling and provide preclinical support for targeting clock activity to improve muscle repair in muscular dystrophy.

## Materials and Methods

### Animals

Mice were maintained in the City of Hope vivarium under a constant 12:12 light dark cycle, with lights on at 6:00 AM (ZT0). All experiments were approved by the IACUC committee of the Beckman Research Institute of City of Hope and carried out in concordance with the IACUC approval. Mice with Tamoxifen-inducible Bmal1 overexpression in satellite cells were generated by crossing Pax7-CreERT transgenic mice (*Pax7^tm1(cre/ERT2)Gaka^*/J, Jackson Laboratory Strain: 017763) with a floxed conditional Bmal1 overexpression allele. Conditional ROSA-26 Bmal1 knock-in allele was generated via CRISPR-mediated gene targeting*(Xiong, Li et al., 2022).* Bmal1-null (Arntltm1Bra, Stock No: 009100) and mdx mutant mice (Dmd-mdx, Stock No: 001801) were originally purchased from the Jackson Laboratory. Mice were crossed for three generations to obtain double homozygote mutants that were used for primary myoblasts isolation.

### Cardiotoxin injury and drug delivery

Acute injury was performed by intramuscular injection (IM) of cardiotoxin (CTX, Sigma-Aldrich 217503) into the tibialis anterior (TA), as previously described(Chatterjee et al., 2019, Chatterjee et al., 2015). Briefly, a 60 μL total volume of 10 μM CTX was injected into 3 sites along longitudinal length of TA muscle. 50uL of 2μM Chlorhexidine and CM002, or PBS as control, were injected into TA after injury in C57BL/6 or mdx mice. Injections were performed at 3:00 PM (ZT9) and tissues were collected in the days post injection (dpi) at 1:00 PM (ZT7).

### Primary myoblast isolation and differentiation

Satellite cells were isolated from pooled hind limb muscle of 8-week-old mice of indicated genotypes using MACS isolation(Motohashi, Asakura et al., 2014) and expanded in culture for primary myoblasts. Briefly, pooled hind limb muscles were collected, finely minced and subjected to collagenase digestion to collect mononuclear cells devoid of multinucleated myofiber. The mononuclear cell digest was washed and seeded on collagen-coated plates, using serial pre-plating to deplete fibroblasts and expand the myoblast population. Myoblasts were subsequently expanded in growth media for 6 passages. Purity of myoblasts obtained were confirmed by uniform differentiation into myosin heavy chain (MyHC)-positive myotubes. Proliferative myoblast cultures were maintained in F-10 medium supplemented with 20% FBS with 2.5 ng/ml bFGF. 2% horse serum (HS)-supplemented DMEM was used for induction of differentiation, as previously described (Kiperman, Li et al., 2023a).

### Myoblasts immunofluorescence staining

C2C12 myoblasts (ATCC CRL-1772), Flox control and sBMKI primary myoblasts, or mdx and human DMD primary myoblasts (Creative Bioarray, CSC-C3604) were seeded at 2 × 10^5^/well onto 6-well plates and maintained in growth media until 80% confluency. DMEM with 2% HS was used to induce differentiation together with treatment of the indicated compounds. Myotubes were fixed with 10% neutral-buffered formalin (Sigma, HT501128-4L) on the days indicated after differentiation, permeabilized and blocked with 1% BSA. Primary antibodies were incubated overnight at 4°C, and secondary antibodies applied at room temperature for 1 h. 4′,6′-diamidine-2′-phenylindole dihydrochloride (DAPI, 1 ug/ml) was used to label nuclei. Images were acquired using an Echo Revolve fluorescence microscope. Primary and secondary antibodies used and the dilutions are described in Supplemental Table 1.

### EdU proliferation assay

To quantify the proliferation of myoblast, cells were seeded at 0.5×10^5^/well in 12-well plates and incorporation of 10 μM of 5-ethynyl-2′-deoxyuridine (EdU) was analyzed following addition for 4 to 6 hours before fixation. For *in vivo* analysis, mice were injected intraperitoneally (IP) of 100 μL EdU overnight prior to tissue collection. Detection of EdU was performed using Click-iT EdU Imaging Kit with Alexa Fluor 488 (Invitrogen). DAPI (1 μg/ml) was used for labeling of nuclei. The total number of EdU^+^ cells was determined using 10 representative fields, and the rate of proliferation was calculated as percentage of EdU^+^/DAPI.

### Skeletal muscle immunofluorescence staining

Muscles were collected and fixed in with10% neutral-buffered formalin (Sigma, HT501128-4L) for 24 hours, washed in PBS and incubated in 30% sucrose solution (Fisher Chemical S5-500) for 48 hours prior to embedding in OCT (Sakura Finetek, 25608-930). 10μm cryosections were collected using TA middle portion, fixed with acetone, and permeabilized with TBST containing 0.4% Tween 20. Endogenous IgG was blocked using non-specific mouse IgG (M.O.M Immunodetection kit, VECTOR Laboratories), with further blocking using 5% goat serum (Vector laboratories S-1000) prior to primary antibody incubation at 4^0^C overnight. Secondary antibodies were applied at room temperature for 1 hour and mounted with ProLong Gold antifade reagent with DAPI (Invitrogen P36931). Muscle cross section area (CSA) was calculated from a total of four representative 10X fields from each muscle and plotted for muscle diameter size distribution.

### Hematoxylin and eosin histology

Muscles were collected and fixed with 4% formaldehyde for 72 hours, washed in 70% alcohol prior to embedding. 10μm paraffin-embedded muscle cross sections were processed for deparaffinization, rehydrated, and stained with hematoxylin and eosin.

### Immunoblot analysis

Total protein (20-40 µg) were extracted using immunoprecipitation lysis buffer (3% NaCl, 5% Tris–HCl,10% Glycerol, 0.5% Triton X-10) and resolved on 10% SDS-PAGE gels followed by western blotting on Immuno-Blot PVDF membranes (Bio-Rad). Primary antibodies were incubated overnight at 4^0^C and secondary antibodies applied at room temperature for 1 hour. Blots were developed using Super Signal West Pico PLUS Stable Peroxide Solution (Thermo Scientific 1863095) and Enhancer Solution (Thermo Scientific CN: 1863094), followed by imaging using a chemiluminescence GE BioSciences AB Amersham Imager 680. Primary and secondary antibodies used and the dilutions are described in Supplemental Table 2.

### RNA extraction and RT-qPCR analysis

Trizol (Invitrogen) or RNeasy miniprep kits (Qiagen) were used to isolate total RNA from snap-frozen tissues or cells. Reverse transcription was carried out using a Revert-Aid RT Reverse Transcription Kit (Thermo Fisher K1691) and ran on a SimpliAmp Thermal Cycler (Thermo Fisher A24811). Quantitative PCR was performed using SYBR Green Master Mix (Thermo Fisher A25742) and ViiA 7 Real-Time PCR System (Applied Biosystems). Relative expression levels of target genes were determined using the comparative Ct method and normalized to 36B4 as internal control. Primer sequences are listed in Supplemental Table 3.

### RNA-seq analysis

RNA-seq reads were aligned with STAR software to the mouse reference genome mm10, and unique reads were quantified using HTSeq-count with GENCODE annotations. The RNA-seq reads were normalized and log-transformed using limma edge R packages. *P*-values and the false discovery rate (FDR) were calculated from raw counts using DESeq2. Fold-change > 1.5, 50% fragments per kilobase of transcript per million mapped reads (FPKM) >0.1, unadjusted *P*-value <0.05, and FDR < 0.25 were used as cut-off for differentially expressed genes. Global analysis heatmaps were produced using heatmap.3 and the gplots package in R, using log2 (FPKM + 0.1) values. The Pearson dissimilarity (1–Pearson correlation coefficient) was used as the distance metric for hierarchical clustering of rows and columns. Expression was centered so that each gene had a mean of 0, and centered expression was capped at –3 and 3, for clustering and visualization.

### Analysis of single-cell RNA sequencing datasets

Published scRNA-seq datasets of uninjured and regenerating mouse skeletal muscle spanning 7 days post injury (dpi)(De Micheli, Laurilliard et al., 2020, Walter, Orton et al., 2024), including GSE143437, GSE159500, GSE162172 and GSE232106 were analyzed in using Scanpy (v1.11.5, Python 3.11). Cell-type annotations, myogenic cell-state labels, MuSCs, non-cycling/cycling/committing MPCs, fusing myocytes and myonuclei, were applied as defined by the original authors. The same methodologies were applied for pseudotime ordering and cell-cycle assignments(De Micheli et al., 2020, Walter et al., 2024). A focused differential-expression comparison of non-cycling versus cycling MPCs was performed using Wilcoxon test. Arntl (Bmal1), CLOCK, Notch and Wnt module scores were computed per cell with Scanpy by applying the Tirosh control-gene-bin method. Relative scores were calculated within this dataset and summarized as violin plots across the ordered cell states, and then directly compared between non-cycling and cycling MPCs by two-sided Mann-Whitney U test with Bonferroni correction. The Notch nodule was comprised of these genes: *Notch1, Notch2, Notch3, Jag2, Dll1, Hes1, Hes5, Hes6, Hey1, Hey2, Rbpj and Nrarp*. The canonical Wnt module included *Ctnnb1, Axin2, Tcf7, Lef1* and *Ccnd1* genes.

### Chromatin Immunoprecipitation (ChIP) analysis

C2C12 cells were collected, crosslinked and sonicated to shear the chromatin according to manufacturer’s instructions of the Simple ChIP Plus Sonication Kit (Cell Signaling Technology 56383). Immunoprecipitation was performed with 5μg Bmal1 antibody (AB93806; Abcam) or rabbit IgG. The immunoprecipitated chromatin fragments were processed and purified using the Simple ChIP Plus Sonication Kit and DNA Purification Buffers (Cell Signaling Technology 14209). Real-time PCR using Taq Pro Universal SYBR qPCR Master Mix (Vazyme Q712-03) was carried out with specific primers flanking the identified E-box sites using genomic primers for Tbp promoter as negative control. Primer sequences are listed in Suppl. Table 4. Data are expressed as fold enrichment over IgG after normalization to 1% input.

### Statistical analysis

Experiments results were analyzed and graphed using PRISM, and data were presented as mean + SD. Each experiment was repeated at least twice. Biological replicates were indicated for each experiment in figure legends. Unpaired two-tailed Student’s t test was used for two group comparisons, and one-way ANOVA for multiple comparisons with Dunnets’s or Tukey’s post hoc analysis were performed as appropriate. P<0.05 was considered statistically significant. GraphPad Prism software (version 6.01) was used for generating graphs, and BioRender was used for schematic illustrations.

## Results

### Circadian clock marks cycling myogenic precursor cells in muscle regeneration and exerts direct transcriptional control of the Notch signaling pathway

The regenerative response induced by acute muscle injury is an established paradigm to study muscle stem cell functions(Charge & Rudnicki, 2004). To explore potential dynamic regulations of clock activity during muscle regeneration, we surveyed clock genes using publicly available sc-RNA-seq datasets that delineated myogenic progression in response to muscle damage from uninjured state to specific days post injury (dpi) from day 1 to 7(De Micheli et al., 2020, Walter et al., 2024). The expression of all clock genes surveyed in cell types identified in muscle regeneration were plotted (**Fig. S1A**), along with their distribution in subgroups within the MuSCs and myogenic progenitor cell populations (MPC, **Fig. S1B**). Interestingly, though neither Bmal1 nor CLOCK was robustly expressed in mature myonuclei, they were both enriched in MuSCs (**Fig. S1B**). Notably, these transcriptional activators of core clock oscillation displayed strong enrichment among subpopulations of cycling and committing vs non-cycling MPCs (**Fig. 1A**). A Pseudotime analysis was applied to trace the lineage trajectory of MuSC from non-cycling to cycling, committing, and fusing MPCs using originally defined cell state annotations based on marker gene expression (**Fig. S2B**) (Walter et al., 2024). Notably, Bmal1 and CLOCK expression displayed a regulatory dynamic along regenerative myogenesis that were maintained at markedly elevated levels in MuCSs with a sharp return to basal suppressed states during lineage commitment and differentiation (**Fig. 1B**). This dynamic regulation aligned closely with Notch gene expression score, a key signaling pathway in determining MuSC proliferative expansion and renewal. We therefore determined the distribution of Bmal1 and CLOCK together with key signals driving MuSC behavior in regeneration (**Fig. S2**). Their aggregate expression among MuSCs steadily increased from day 2 of regeneration (**Fig. S2A**). Among the distinct myogenic subclusters (**Fig. S2B & S2C**), the levels of both core clock regulators were induced in MuSC and cycling MPCs, with gradual dampening of expression along commitment and differentiation of MPCs (**Fig. 1C & S2D**). Notably, this regulation mirrored that of the composite Notch pathway score, while partially correlated with Wnt score among MuSC and MPCs, though Wnt was also maintained at high levels in mature myonuclei. The dynamic clock induction in MPCs actively participating in regeneration closely aligned with Notch activation, suggesting its potential involvement in this signaling event.

**Figure 1.**
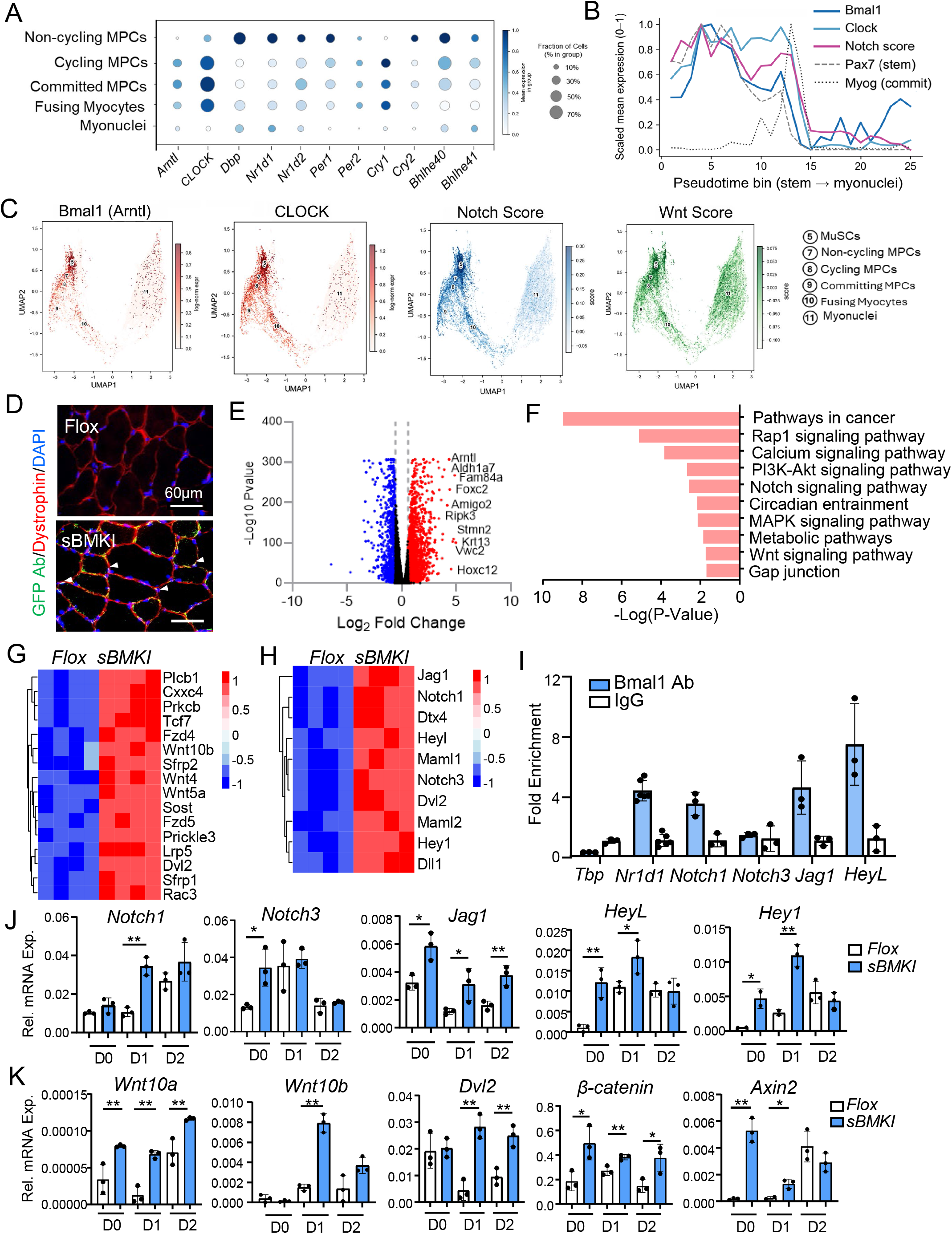
Circadian clock direct transcriptional control and functional regulation of the Notch and signaling pathway. (A-C) Identification of molecular clock activation in muscle stem cell populations from sc-Seq analysis during muscle regeneration. Published scRNA-seq datasets GSE143437, GSE159500 and GSE162172 and GSE232106 were used in the analysis. (A) Dot plot analysis of core clock and clock output gene expression across ordered myogenic states among non-cycling, cycling and committed MPCs, fusing myocytes, and myonuclei. Dot size denotes the fraction of expressing cells within each group; color gradient denotes scaled mean expression. (B) Pseudotime trajectory analysis across the ordered myogenic states. MPCs were plotted along the pseudotime lineage trajectory based on published results that grouped into 25 bins across the ordered myogenic states spanning stem cells to myonuclei. Scaled mean expression of Bmal1, Clock, and the computed Notch signaling score (see Methods) is shown together with the MuSC marker Pax7 (dashed line) and committed MPC marker Myogenin (dotted line) across binned pseudotime trajectory. (C) UMAP feature plots of the myogenic compartment showing Bmal1 and CLOCK enrichment within MPCs subgroups along lineage progression in regeneration with correlation of Notch and canonical Wnt signaling scores. (D) Representative immunofluorescence staining using GFP antibody (green) with Dystrophin (red) of Flox control and sBMKI TA muscle. Scale bar: 60 μm. Arrowheads denote satellite cells with GFP expression. (E, F) Volcano plot representation of differentially expressed genes (E), and KEGG pathway enrichment analysis (F) of isolated sBMKI primary myoblasts as compared to Flox controls (n=3/group). (G, H) Heatmap of significantly expressed genes in Flox and sBMKI primary myoblast involved in the Notch (E), or Wnt (F), signaling pathways. (I) ChIP-qPCR analysis of chromatin occupancy of Bmal1 on identified E-box sites of proximal promoter regions (+2Kb from TSS) of genes involved in Notch signaling. E-box site on Nr1d1 gene was used as a positive control. N=3-6/group. (J, K) RT-qPCR analysis of dynamic expression of components of Notch (H), and Wnt (I) pathways in in Flox and sBMKI primary myoblast before and following myogenic differentiation for 2 days. N=3/group. * and **: p<0.05 or 0.01 by Student’s t test.

To experimentally determine clock activation of stem cell functions in skeletal muscle, we established a clock gain-of-function model with tamoxifen-inducible Bmal1 conditional overexpression in satellite cells (sBMKI). The was achieved by crossing the Flox Bmal1 allele (Flox), containing Bmal1 Flag-tagged cDNA with bicistronic GFP flanked by a Flox-Stop-Flox sequence, with the Pax7-CreERT2(Murphy, Lawson et al., 2011) mice(Xiong et al., 2022). Five days following tamoxifen induction, Flag-tagged Bmal1 expression was detected in skeletal muscle groups examined together with GFP, but not in the liver or heart (**Fig. S3A**). The ectopic Bmal1 overexpression was detected in satellite cells residing outside the sarcolemma as shown by GFP antibody with dystrophin co-staining (**Fig. 1D**). We first interrogated clock-controlled outputs in modulating satellite cell properties through transcriptomics profiling using satellite cell-derived primary myoblasts obtained from sBMKI and Flox controls. Global analysis of differentially expressed genes revealed up-regulation of 2659 genes in sBMKI myoblasts comprising 16.6% of all transcripts detected, as compared to 1022 (6.4%) displaying down-regulation (**Fig. 1E**), as expected of Bmal1 gain-of-function in these cells. Bmal1 (Arntl) was among the top up-regulated genes with a ∼4.4-fold induction. As shown by KEGG pathway analysis, known clock-modulated processes, including cancer-related, circadian entrainment and metabolic regulations(Miller et al., 2007), were among the top up-regulated biological processes (**Fig. 1F**). Notch and Wnt signaling are major mechanisms driving MPC myogenic lineage progression in muscle regeneration(Brack et al., 2008, Conboy & Rando, 2002). Notably, both pathways were found to be among the top enriched processes (**Fig. 1F**), with their key signaling components markedly induced (**Fig. 1G & 1H**). These findings implicated clock transcriptional control of Wnt and Notch pathways in MuSCs that may impact their activation and myogenic properties. Specific components of the Wnt signaling cascade were previously identified as direct Bmal1 targets in myogenic progenitors(Chatterjee et al., 2013, Guo, Chatterjee et al., 2012). To dissect whether genes involved in Notch signaling could also be under the direct transcriptional control of clock, we surveyed E-box elements that meditate Bmal1 chromatin binding within the proximal promoter regions of candidate targets(Rey, Cesbron et al., 2011).

Using the known Bmal1 target *Nr1d1* promoter as a positive control, ChIP-qPCR analysis revealed that among these candidates, *Notch1*, *Jagged 1* (*Jag1*) and *HeyL* promoters displayed Bmal1 occupancy on identified E-boxes (**Fig. 1G**), while *Notch3* showed minimal binding affinity. Furthermore, when compared with Flox controls during normal proliferative growth or myogenic differentiation, these Notch signaling genes, *Notch 1*, *Jag1* and *HeyL*, were robustly induced in sBMKI myoblasts (**Fig. 1J**). Importantly, most Notch signaling components examined in sBMKI cells were modulated similarly as controls following day 2 of myogenic differentiation, suggesting a normal response to myogenic stimuli. Similar up-regulations of Wnt pathways genes that were identified as Bmal1 targets(Chatterjee et al., 2013), including *Wnt10a*, *Wnt10b*, *Dvl2* and *β-catenin*, were found in sBMKI myoblasts under these conditions (**Fig. 1K**). Notably, *Axin2* expression level, indicative of the transcriptional activity of the Wnt cascade, was robustly up-regulated early during differentiation, though it was repressed to a comparable level as that of the controls on day 2.

Under normal conditions without injury, the overexpression of Bmal1 in satellite cells did not elicit significant impact on normal muscle growth or function. The wet weight of distinct muscle groups, Tibialis Anterior (TA) and gastrocnemius (GN), were comparable with Flox controls (**Fig. S3B & S3C**). Assessment of myofiber cross-section area *via* H/E histology (**Fig. S3D**) or laminin staining (**Fig. S1E**) revealed comparable fiber sizes. Muscle function as determined by grip strength were also similar between these groups (**Fig. S3F**).

### Genetic clock gain-of-function in satellite cells prolonged the myogenic phase with a delayed peak in muscle regeneration

In response to tissue damage, regeneration of the skeletal muscle follows a time course driven by proliferative expansion of satellite cells upon activation and subsequent myogenic lineage progression to generate nascent myofibers and fusing to form mature myocytes(Charge & Rudnicki, 2004). To determine the functional properties of MuSCs with ectopic expression of Bmal1, the sBMKI mice were subjected to acute injury by cardiotoxin and the dynamic regenerative response were examined for 30 days post injury till dpi 30. Acute injury in the TA muscle elicited a strong response of circadian clock and regenerative myogenesis by MuSC clock gain-of-function. As expected, Bmal1 protein was elevated at basal state without injury in sBMKI TA muscle (**Fig. 2A**). Both Bmal1 and CLOCK protein steadily rose following injury along the time course of regeneration, with marked inductions at dpi 5 to 30 (**Fig. 2A**). In contrast to the endogenous Bmal1 protein that was induced during regeneration peaking at dpi 3-5 followed by a return to basal levels at day 30, the high levels of Bmal1 and CLOCK protein induction were likely due to the formation of new myofibers containing ectopic Bmal1 at later stages of regeneration. Further analysis revealed an interesting regulatory dynamic of the myogenic response observed in the sBMKI mice (**Fig. 2B**). At the early regenerative stage of 3 days after injury, the myogenic response, as indicated by the induction of *Pax7*, *Myf5*, *Myod1* and *Myogenin* gene expression, displayed a modest attenuated tendency in sBMKI mice as compared to controls. In contrast, at later time points of regeneration from dpi 5 to 14, their expression levels were mostly elevated. At dpi 30 when regeneration was completed, the expression of these genes in sBMKI mice returned to comparable levels as in the Flox controls. Examination of the protein levels of myogenic regulators further validated the gene expression analysis (**Fig. 2C**). The reductions of Pax7, Myf5 and Myogenin protein were evident at dpi3 and 5 in sBMKI mice. However, at later stages of regeneration of dpi 14 as well as dpi30, when the early induction of these factors returned to undetectable levels in the controls, as expected of normal regeneration, these proteins remained elevated in sBMKI mice. These findings revealed a distinct regenerative trajectory in sBMKI mice, displaying an attenuated myogenic induction at early stage of regeneration with a prolonged myogenic drive beyond the normal response, suggesting a delayed but augmented total regenerative phase in mice with MuSC clock overexpression. Laminin immunostaining was used to examine nascent myofiber development via determination of myofiber size progression during regeneration. As shown by cross-section area distribution at dpi5 in sBMKI mice, there was a marked shift of myofibers toward 500µm^2^ or smaller sizes as compared to controls, indicative of nascent regenerated fibers at this early stage of regeneration (**Fig. 2D**). In comparison, by dpi 14, a reduction of myofibers in the small size range was evident in sBMKI mice, together with a tendency toward shifting to larger size range (**Fig. 2E**). At dpi 30 when the normal regeneration time course was largely complete in the controls, sBMKI TA displayed a reduction of myofibers at the smaller end of the size distribution (500-1,000 µm^2^), while the abundance of larger fibers of ≥2,000 µm^2^ diameters were significantly elevated (**Fig. 2F**). These analyses together indicated that clock gain-of-function in MuSCs resulted in a delayed early regenerative response, though the regenerative phase was prolonged with an extended timeline.

**Figure 2.**
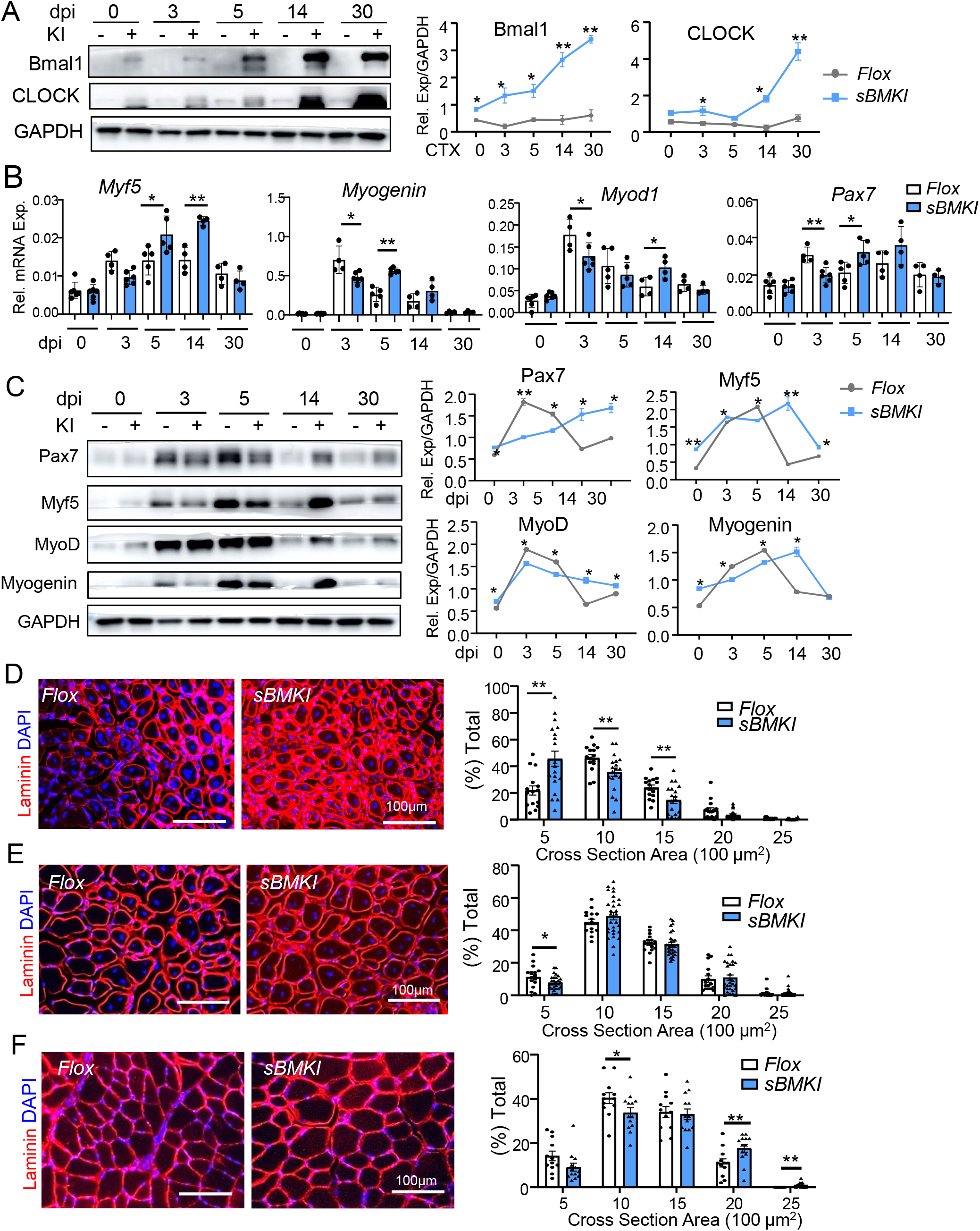
Satellite cell clock gain-of-function prolongs the regenerative myogenic phase of muscle regeneration. (A-C) Gene expression analysis of the dynamic myogenic response along the regeneration time course following cardiotoxin injury of TA muscle. Representative immunoblot analysis of clock protein induction (A), myogenic response (C), and RT-qPCR analysis (B) of Flox and sBMKI mice at indicated times till 30 days after injury. dpi: days post injury. For immunoblot analysis, each lane represents a pooled sample of 5-7 mice/group, with quantification of three repeats. (D-F) Representative images of laminin immunofluorescence staining of myofiber cross sections at dpi 5 (D), dpi 14 (E) and dpi 30 (F) with corresponding quantitative analysis of myofiber section area. Scale bar: 100μm.

### Clock gain-of-function in MuSCs promotes satellite cell proliferative expansion and myogenic progenitor properties

Using immunofluorescence staining of eMyHC to identify nascent myofibers, we next examined the delayed regeneration phase in sBMKI mice. Despite an attenuated myogenic response early in regeneration (**Fig. 2 & Fig. S4**), newly regenerated eMyHC^+^ myofibers at dpi 5, beyond the normal peak of new myofiber generation at dpi 3, were more abundant in sBMKI mice, suggesting a delayed yet augmented peak of nascent myofiber formation in line with the dynamics of myogenic response (**Fig. 3A**). EdU labeling of proliferative MuSCs with Pax7 co-staining revealed that the rate of satellite cell proliferation detected at day 7 dpi, near the end of normal expansion phase, were significantly higher in sBMKI mice, indicative of enhanced satellite cell proliferative expansion (**Fig. 3B**). A temporal progression from Notch pathway activation at early regeneration to Wnt signaling at later stages is required for normal regenerative myogenesis(Brack et al., 2008). Given clock transcriptional regulation of Wnt signaling components(Chatterjee et al., 2013, Guo et al., 2012) and current findings of Notch pathway genes as direct Bmal1 targets, the transcriptional control by clock may temporally coordinate Notch and Wnt signaling leading to prolonged regenerative response in sBMKI mice. Analysis of both pathways along the regeneration time course revealed that a major Notch receptor in MuSCs, *Notch1*, and Notch signaling downstream effector *HeyL* were induced at early stages of regeneration, though both were effectively suppressed at the completion of regeneration at dpi 30 similarly as the controls (**Fig. 3C**). In contrast, a known Bmal1 target within the Wnt pathway, *Wnt10a*, was induced at day 5 and 14 of regeneration. Expression of the Wnt receptor *Fzd5* was attenuated at dpi 3 but robustly elevated at dpi 14 that persisted till day 30 (**Fig. 3C**). Thus, forced clock expression in MuSCs induced Notch activity early in regeneration while prolonging Wnt pathway up-regulation later from dpi 5 till 30, revealing a sequential switch from activating Notch to Wnt signaling during the progression of myogenesis. Given these pathways in regulating MuSC self-renewal, proliferation and lineage commitment in muscle regeneration, we isolated satellite cells on day 5 after CTX injury and determined their activation and myogenic commitment properties (**Fig. 3D & 3E**). Identification of proliferative satellite cells via EdU/Pax7 labelling revealed a ∼27% increase of proliferative rate among these cells isolated from sBMKI mice as compared to that of controls (**Fig. 3D**). Furthermore, a significantly higher proportion of cells double positive for MyoD and Myf5 indicative of activated myogenic progenitors(Kuang et al., 2007) were found in the satellite cells from sBMKI mice (**Fig. 3E**).

**Figure 3.**
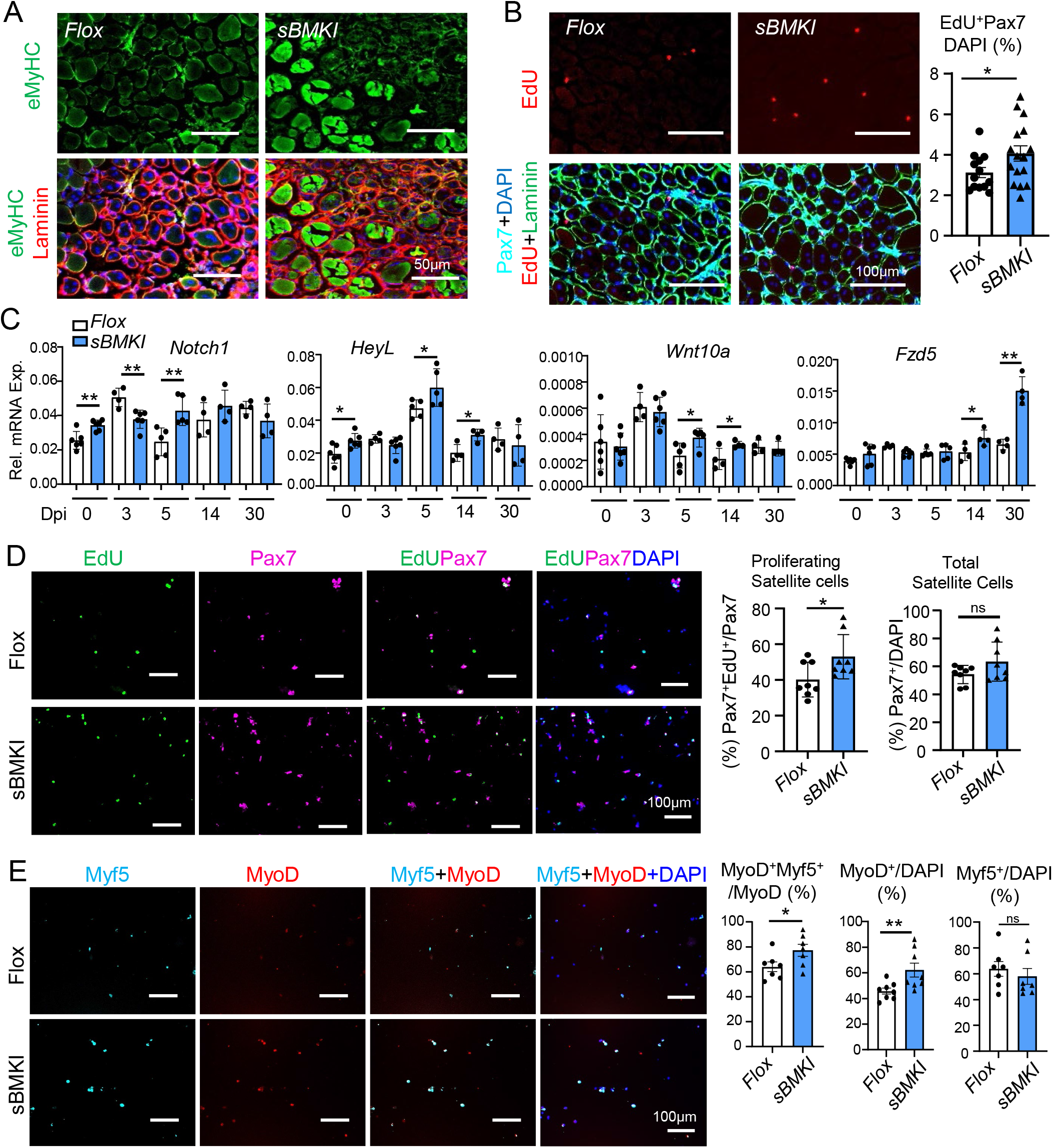
Clock promotes satellite cell proliferative expansion and myogenic progenitor function. (A) Representative images of immunofluorescence staining of myosin heavy chain (MyHC, green) and laminin (red) to assess nascent myofiber formation in Flox and sBMKI mice at dpi5. (B) Representative images of Pax7 immunofluorescence staining (green) with EdU fluorescence (red) and the quantitative analysis of proliferating satellite cells in TA muscle of Flox and sBMKI mice 7 days after CTX injury. (C) RT-qPCR analysis (B) of Notch and Wnt pathway genes in Flox control and sBMKI mice at indicated times along the regeneration time course. N=5-6 mice/group. * and **: p<0.05 or 0.01 by Student’s t test. (D & E) Representative images of immunofluorescence staining of Pax7 (ed) with EdU (Green) fluorescence labeling to assess proliferation of satellite cells (D), and MyoD (red) or Myf5 (Cyan) to assess MuSC properties (E) using satellite cells isolated from Flox and sBMKI mice at dpi 5 after CTX injury (n=5/group). Scale bar: 100μm.

Furthermore, we explored whether enhanced Notch signaling mediated the induction of MuSC proliferative expansion by Bmal1 overexpression, using primary myoblasts derived from satellite cells via *ex vivo* expansion. Under the normal proliferative growth condition, satellite cell-derived sBMKI myoblasts displayed a significant ∼38% increase in proliferation than that of the controls (**Fig. 4A**). Following the induction of myogenic differentiation using 2% horse serum, while the proliferation of Flox controls dropped as expected, elevated level of proliferative rate of sBMKI cells was maintained, in line with the observation of prolonged myogenic response *in vivo*. Further analysis of cell cycle regulators revealed induction of *Cyclin D1* and *Cdk4* genes in proliferative sBMKI myoblasts, with their expression similarly inhibited as the controls following differentiation (**Fig. 4B**). Interestingly, despite the up-regulation of cell cycle regulators, expression of myogenic markers, *Myod1*, *Myogenin* and *eMyHC,* were markedly elevated in these cells than that of *Flox* controls, suggesting enhanced myogenic differentiation (**Fig. 4C**). To explore whether the increased proliferation of sBMKI myoblasts was mediated by Notch signaling, we tested a specific inhibitor CB103 that blocks the transcriptional activation of Notch pathway(Kiperman, Li et al., 2023b). A low concentration of CB103 (2uM) was sufficient to abolish the increased proliferative rate of sBMKI myoblasts to a level comparable to that of controls (**Fig. 4D & 4E**), suggesting that Notch activity indeed underlies the proliferative phenotype induced by Bmal1 overexpression.

**Figure 4.**
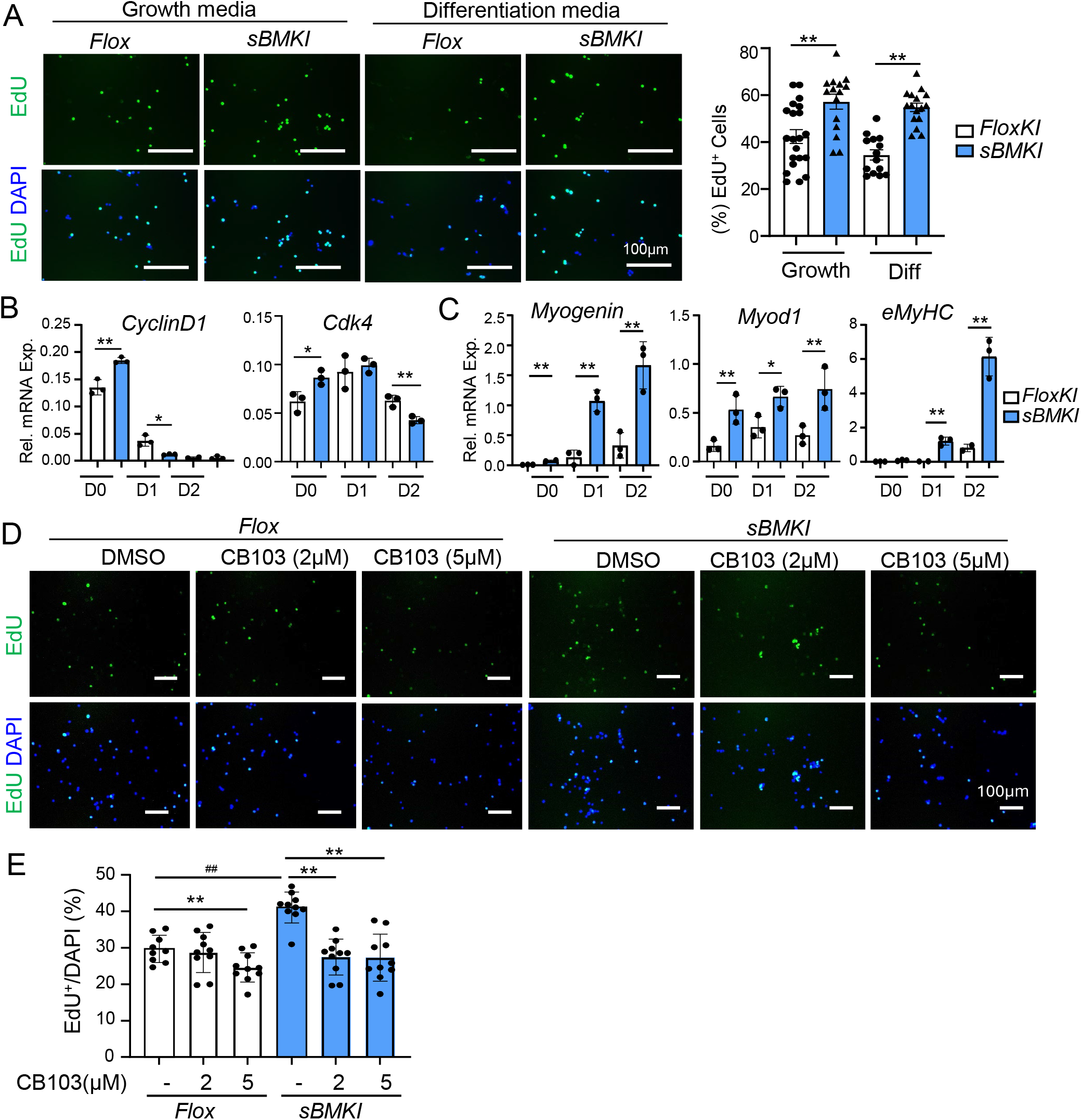
Cell-autonomous effect of Bmal1 on promoting myogenic progenitor proliferation. (A) Representative images of EdU staining with quantitative analysis of Flox and sBMKI primary myoblasts under a proliferative growth condition and following one day of differentiation in 2% horse serum-supplemented media. (B, C) RT-qPCR analysis of dynamic expression of components of cell cycle regulators (B) and myogenic factors (C) in Flox and sBMKI primary myoblasts before and during myogenic differentiation for 2 days. N=3/group. * and **: p<0.05 or 0.01 by Student’s t test. (D, E) Representative images of EdU staining (D) with quantitative analysis (E) to determine effect of the Notch inhibitor CB103 on blocking Flox and sBMKI primary myoblasts proliferation under the proliferative growth condition. *, **: p<0.05 or 0.01 CB103 vs. DMSO control, and ##: Flox control vs. sBMKI.by one-way ANOVA with Dunnet’s post-hoc analysis.

### Pharmacological activation of clock by chlorhexidine and CM002 promotes myogenesis via Notch and Wnt signaling

We recently identified a clock activator, chlorhexidine (CHX), which displayed strong pro-myogenic properties via activation of Wnt signaling(Kiperman et al., 2023a). Based on the chemical scaffold of this original hit, we performed medicinal chemical modifications and generated a series of analogs. These molecules were subsequently subjected to a Per2::Luc luciferase biochemical screen to determine clock-activating properties followed by a functional screen to assess their pro-myogenic activities. This screening pipeline led to the discovery of a new lead compound, CM002, that displayed increased clock-modulatory activity (**Fig. 5A & S5A**). Viability assay revealed these compounds did not cause cellular toxicity within 0.2-1 μM in myoblasts, and CM002 displayed lower toxicity than CHX at higher concentrations examined (**Fig. S5B**). Consistent with their clock-activating properties, both molecules were sufficient to induce the expression of identified Notch pathway genes along with activation of Wnt signaling as indicated by *Axin 2* level (**Fig. 5B**), with CM002 treatment leading to a modest increase in clock target expression. Consistent with its effect on inducing Notch pathway gene expression, CM002 was sufficient to promote the chromatin occupancy of Bmal1 on identified clock-controlled Notch pathway gene promoters, including *Notch 1*, *HeyL* and *Jag1* (**Fig. 5C**). The fold of induction stimulated by CM002 was promoter-specific, with *Notch 1* highly responsive resulting in a ∼34-fold induction as compared to basal Bmal1 occupancy on this promoter. This finding promoted a further interrogation of the CM002 potential in eliciting circadian time-dependent clock activation of Notch genes. Myoblasts were treated with CM002 at indicated times after synchronized by serum shock to determine its modulation of Notch pathway genes. Circadian oscillation of Bmal1 protein was induced by synchronization, as expected, with its peak abundance occurring between CT11-17 (CT: circadian time, **Fig. S5C**). We thus treated cells at the circadian time window between CT11-17 as compared with its trough period at CT23-29 with determination of induction of Notch pathway genes. This led to a striking finding of the marked inductions of Notch transcripts, including *Notch 1*, *Noth 3* and *Jag 1,* by CM002 when treated at CT11-17 that were significantly higher than the minimal response observed at CT23-29 (**Fig. 5D**), consistent with the heightened clock activity observed at CT11-17. In comparison, expression of *HeyL* revealed a distinct different response, with robust CM002 inductions at both time points. As an effector of Notch signaling, this response of *HeyL* could be potentially due to its function in integrating upstream Notch activity altered by CM002. Consistent with the activity of CHX and CM002 on inducing Notch signaling, both compounds stimulated the proliferation of primary myoblasts, though CM002 appeared more efficacious under this condition (**Fig. 5E & S5D**).

**Figure 5.**
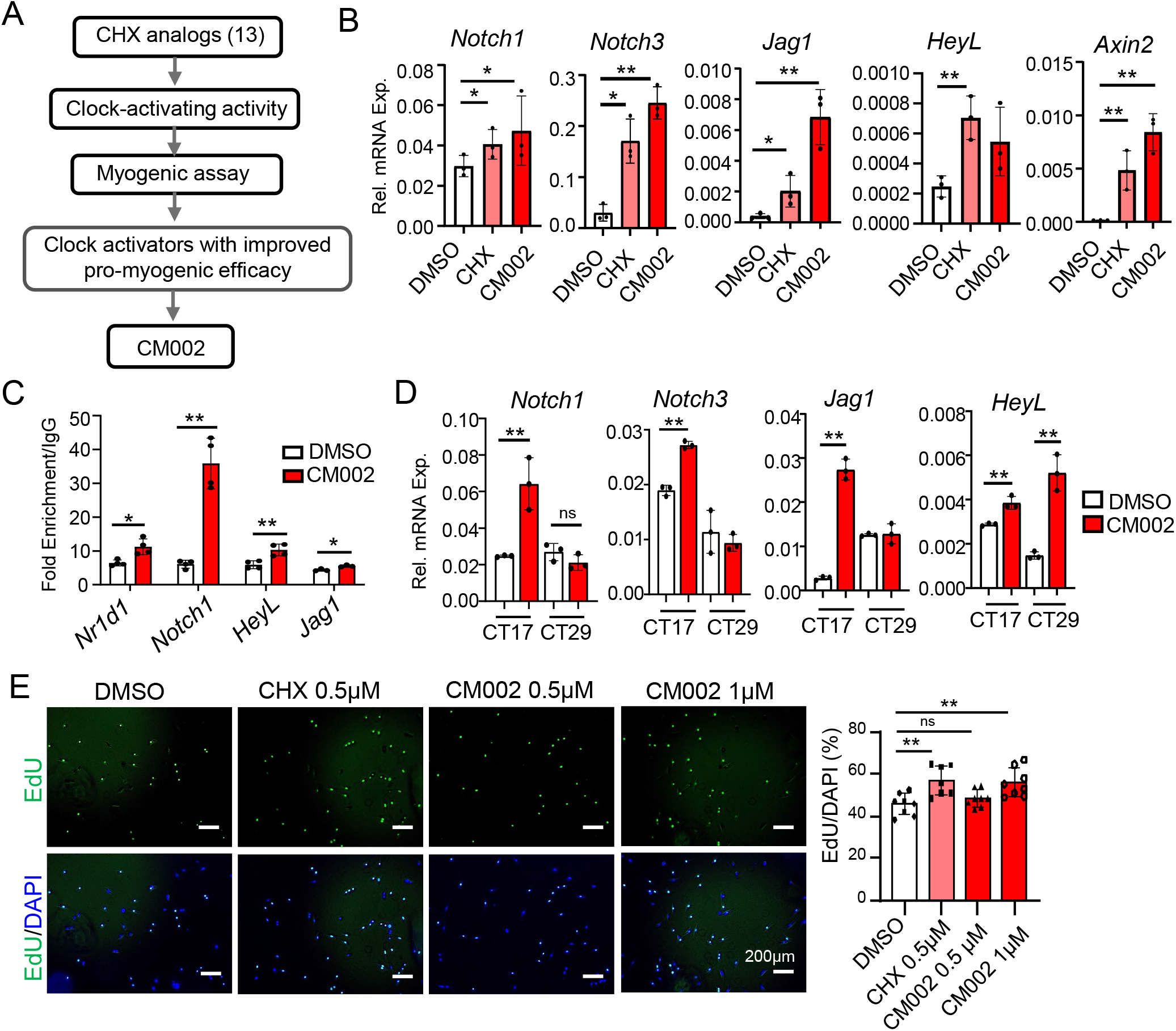
Effect of clock-activating molecules on inducing Notch and Wnt pathway. (A) Schematic flow chart of generating analogs of chlorhexidine leading to the screen and identification of CM002 as a novel clock-activating lead compound. (B) RT-qPCR analysis of Notch and Wnt pathway genes of C2C12 myoblasts treated with CHX (0.5µM) or CM002 (0.5µM) for 6 hours. * and **: p<0.05 or 0.01 CM002 or CHX vs. DMSO control by Student’s t test. (C) ChIP-qPCR analysis of chromatin occupancy of Bmal1 on identified target promoters involved in Notch signaling. C2C2 myoblasts were treated with vehicle (0.1% DMSO) or 0.5µM CM002 for 6 hours prior to chromatin extraction (n=3). * and **: p<0.05 or 0.01 CM002 vs. DMSO control. (D) Effect of time-dependent effect of CM002 on modulating Notch signaling components by RT-qPCR analysis in synchronized myoblasts. Serum shock by one hour treatment of 20% serum was used for synchronization. Cells were treated with DMSO or CM002 (0.5uM) for 6 hours prior to sample collection at CT17 and 29 after serum shock. *, **: p<0.05 or 0.01 CM002 vs. control by one-way ANOVA with Tukey’s post-hoc analysis. (E) EdU fluorescence labeling to assess CHX and CM002 effect on primary myoblast proliferation under normal growth conditions. (n=7-8/group). *, **: p<0.05 or 0.01 CM002 vs. DMSO by one-way ANOVA with Tukey’s post-hoc analysis.

### Clock activation promotes the myogenic differentiation of mouse and human dystrophic myoblasts

Based on the function of clock in promoting myogenesis (Chatterjee et al., 2013), we compared the effects of these clock-activating molecules on myogenic differentiation. Immunofluorescence staining of myosin heavy chain (MyHC), a mature myocyte marker, demonstrated that 0.5 µM CHX markedly enhanced mature myofibers formation after 3 days of differentiation, as compared to DMSO control (**Fig. 6A**). In comparison, MyHC^+^ myofibers treated with 0.5 µM of CM002 were increased than that of CHX with further increase observed at 1 µM, suggesting a stronger efficacy in stimulating myogenic differentiation. When tested in primary myoblasts isolated from mdx mice, a preclinical model for Duchenne Muscular Dystrophy (DMD), CHX was sufficient to promote the differentiation of dystrophin-deficient myoblasts, as indicated by MyHC^+^ myofiber staining 3 days after of myogenic induction (**Fig. 6B**). Furthermore, in Bmal1-deficient mdx primary myoblasts obtained from BMKO/mdx double-null mice(Gao, Xiong et al., 2020), this effect of CHX was completely abolished, indicating that the effect of CHX on promoting myogenesis is clock-dependent. Similar to the higher efficacy of CM002 in promoting myogenesis in C2C12 myoblasts, it demonstrated a stronger effect in primary myoblasts as shown by increased abundance of MyHC^+^ myotubes following differentiation (**Fig. 6C**). To explore the potential for augmenting clock actions to promote regenerative repair applicable to dystrophic diseases, we tested these compounds on proliferation and myogenic differentiation of primary myoblasts derived from DMD patients. In the DMD primary myoblasts, similar effects of CHX and CM002 on inducing proliferation were observed (**Fig. S5D**). In addition, both CHX and CM002 were able to induce up-regulations of Bmal1 protein together with markedly elevated levels of MyoD and Myogenin, suggesting activation of the myogenic response with comparable inductions (**Fig. 6D, 6E & S5C**). Collectively, these findings revealed that pharmacological clock activation is sufficient to enhance myogenesis with potential applications in dystrophic diseases.

**Figure 6.**
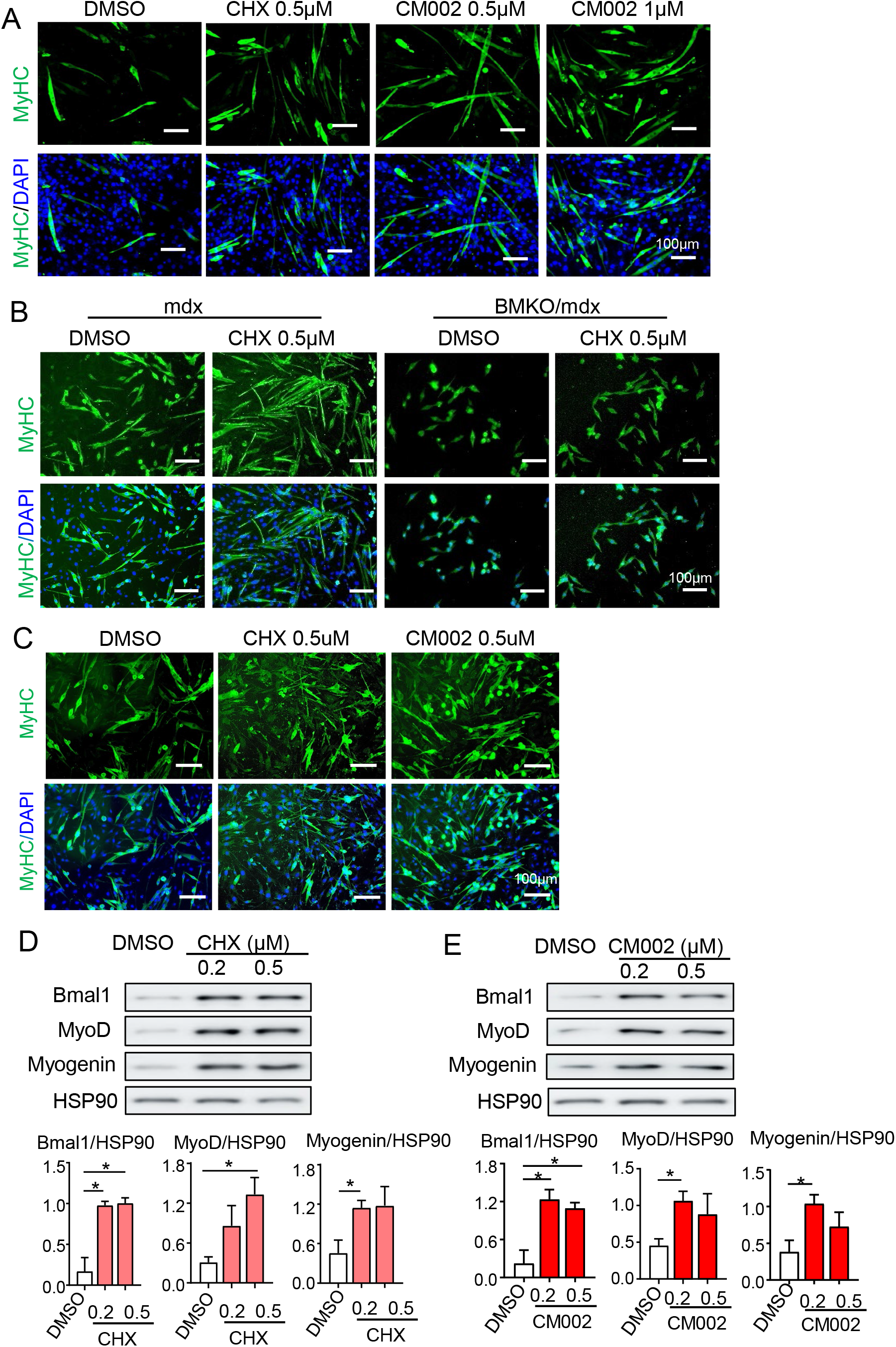
Clock activators promote myogenic differentiation of mouse and human dystrophic myoblasts. (**A**) Representative images of immunofluorescence staining of myosin heavy chain (MyHC, green) of C2C12 myoblast myogenic differentiation at day 4 with chlorhexidine (CHX) or CM002 treatment at indicated concentrations. (B) Representative images of MyHC staining of differentiation of primary myoblasts derived from mdx and Bmal1-deficient mdx mice (BMKO/mdx) treated with DMSO or CHX (0.5 μM). Primary myoblasts were differentiated for 3 days using 2% horse serum. (C) Representative images of MyHC staining of mdx primary myoblasts differentiation for 3 days with 0.5 μM of CHX or CM002 treatment. (D, E) Representative images of immunoblot analysis of the effect of CHX (D) and CM002 (E) on clock and myogenic gene regulation using human primary myoblasts with DMD following 3 days of differentiation. Quantitative analysis is based on three independent repeats. *p<0.05 and ** p<0.01 CM002 or CHX treatment vs. DMSO control.

### Clock activators display pro-myogenic activities upon acute muscle injury and in chronic dystrophic milieu

Given the pro-myogenic effects we demonstrated for these clock-activating molecules, we next tested their potential for enhancing regenerative repair *in vivo* upon acute muscle injury or with chronic damage in the dystrophic disease milieu. Four daily intramuscular deliveries of CHX or CM002 into the TA muscle, administered at one day after cardiotoxin injury, were used to assess their effect on regenerative myogenesis (**Fig. 7A**). eMyHC immunofluorescence staining of regenerated neomyofiber formation was assessed at dpi 5 together with laminin staining (**Fig. 7B**). While vehicle-treated controls displayed low levels of new myofibers as expected at this stage of regeneration, both CHX and CM002-treated regenerating TA demonstrated substantial numbers of nascent myofibers with eMyHC^+^ staining, suggesting sustained new myofiber regeneration. Interestingly, laminin staining revealed a marked increase in the abundance of myofibers observed per section area in CHX or CM002-treated regenerating muscles, indicative of an enhanced regenerative response with increased number of regenerating myofibers. Myofiber size distribution at this stage indeed revealed a robust increase in the abundance of smaller myofibers manifested as a shift toward smaller cross section area (**Fig. 7C**), with a reduction of myofibers within the next group of larger cross section area. Myofibers of medium size ranging from 2000-2500 μm^2^ were increased, suggesting a tendency toward accelerated regenerating myofiber size incretion at advance stages with depletion in the middle range. Myf5, MyoD and Myogenin protein levels were increased in TA muscles treated with CHX or CM002, with a more robust response induced by CM002 than that of CHX (**Fig. 7D & 7E**), though the induction of Pax7 was comparable. Thus, clock-activating molecules are capable of promoting myogenic response *in vivo* with sustained new myofiber formation following acute muscle damage.

**Figure 7.**
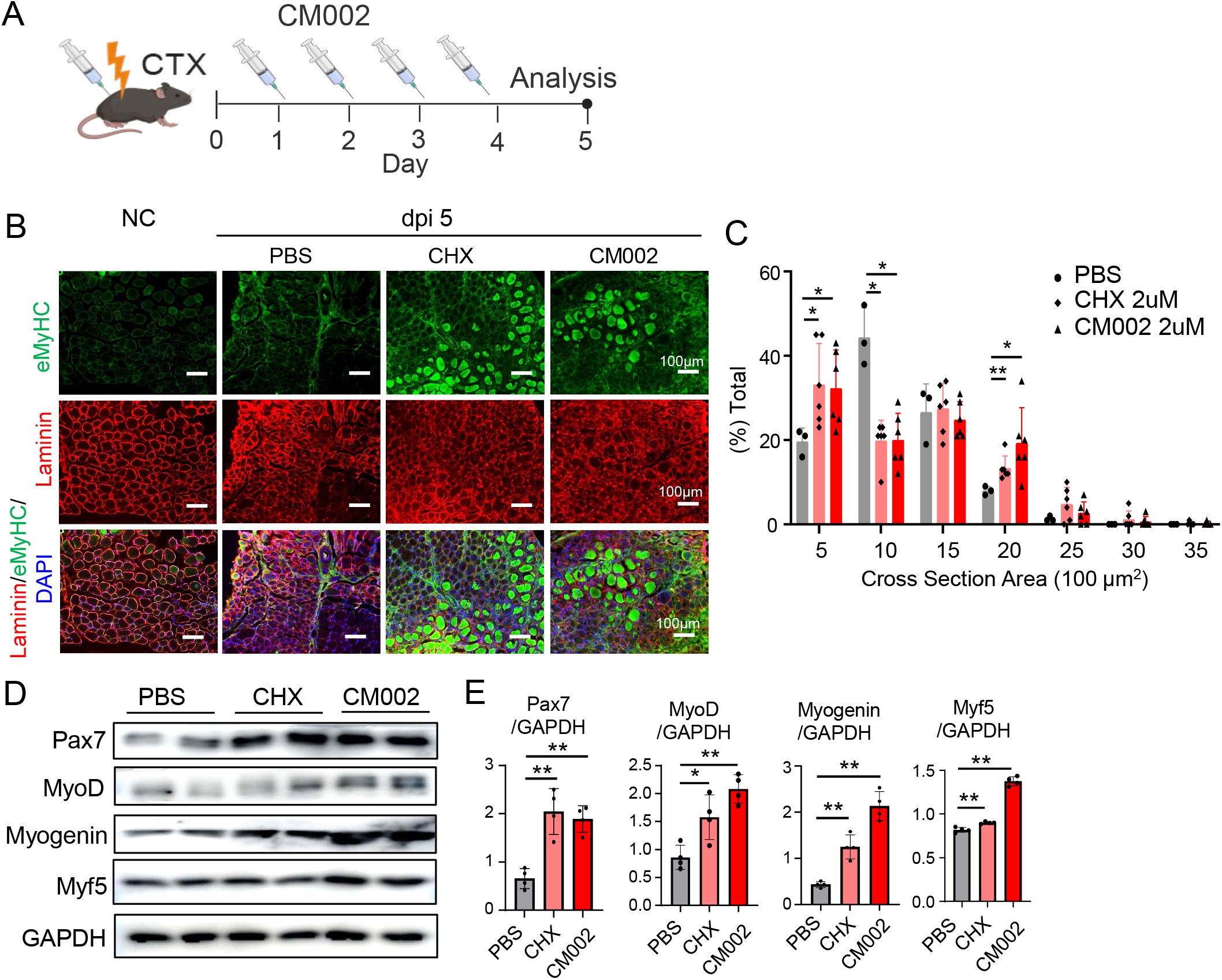
Effect of pharmacological clock activation on regenerative myogenesis upon cardiotoxin-induced injury. (A) Schematic representation of CM002 treatment time course following CTX injury of TA muscle. One day after CTX injury, 50ul of CHX (0.5uM in PBS) or CM002 solution (0.5uM in 0.1% DMSO) was injected into injured TA for 4 days with samples collected at dpi 5 for analysis. (B, C) Representative immunofluorescence staining analysis of embryonic myosin heavy chain (eMyHC) and laminin to examine new myofiber synthesis in TA muscle at dpi5, with quantitative analysis of myofiber cross section area distribution using laminin-stained images (C). (D, E) Immunoblot analysis of genes involved in regenerative myogenesis following injections with CHX or CM002 in CTX-injected TA muscle, with quantitative analysis using three repeats (E). Each lane represents pooled samples of 4-6 mice/group, with quantitative analysis from three repeats.

We next explored whether these effects of CHX and CM002 could be applicable to enhance regeneration in the dystrophic condition with chronic muscle damage, using a similar delivery to the TA muscle of mdx mice(Bulfield, Siller et al., 1984) (**Fig. 8A**). Notably, both compounds elicited marked inductions of key clock regulators, Bmal1, CLOCK and its direct target gene Rev-erbα, in the dystrophic muscle (**Fig. 8B**), with CM002 displaying stronger effect than that of CHX. These molecules also induced robust up-regulations of key signaling components of the Notch and Wnt pathways, as demonstrated for the identified direct Bmal1 targets within these pathways (**Fig. 8C**), indicative of their activation of clock outputs under this condition. Protein levels of cleaved Notch, a critical mediator of intracellular Notch signaling, and the Notch pathway effector, Hes1, were increased by both compounds, further validating enhanced Notch signaling activity in treated mdx muscles (**Fig. 8D**). β-catenin protein was also increased, suggesting induction of Wnt signaling by these compounds. Consistent with their differing efficacy on inducing clock regulators, CM002 effect on these Notch pathway components were stronger than that of CHX. Additional analysis of CHX and CM002-treated muscle revealed elevated Pax7 and Cyclin D1 proteins, along with strong inductions of myogenic factors (Myf5, MyoD & Myogenin, **Fig. 8E**), indicating enhanced regenerative myogenesis in the dystrophic disease milieu by both compounds with comparable efficacy. The proliferative rate of satellite cells in dystrophic muscle following treatments was assessed using EdU labelling (**Fig. 8F**), and we found that compared to vehicle controls, the proportion of EdU^+^/Pax7^+^ satellite cells in CHX and CM002-treated cohorts were increased by ∼52%, while the total number of Pax7^+^ satellite cells were comparable between these groups, suggesting an increase in the proportion of cycling satellite cells without significant expansion of total pool size. Staining of nascent myofibers revealed a strong increase in eMyHC^+^ neo-myofiber formation in CHX or CM002-treated groups (**Fig. 8G**). In contrast to the findings from CTX injury, a striking increase in the abundance of small regenerating myofibers was dominant in both compound-treated groups. This resulted in an overall shift of myofiber size distribution toward smaller myofibers and a corresponding reduction of larger sizes, in line with the emergence of small nascent myofibers detected by eMyHC staining following treatments (**Fig. 8H**). The current findings of the pro-myogenic efficacies of clock-activating molecules in mdx muscle suggest their potential for dystrophic diseases applications.

**Figure 8.**
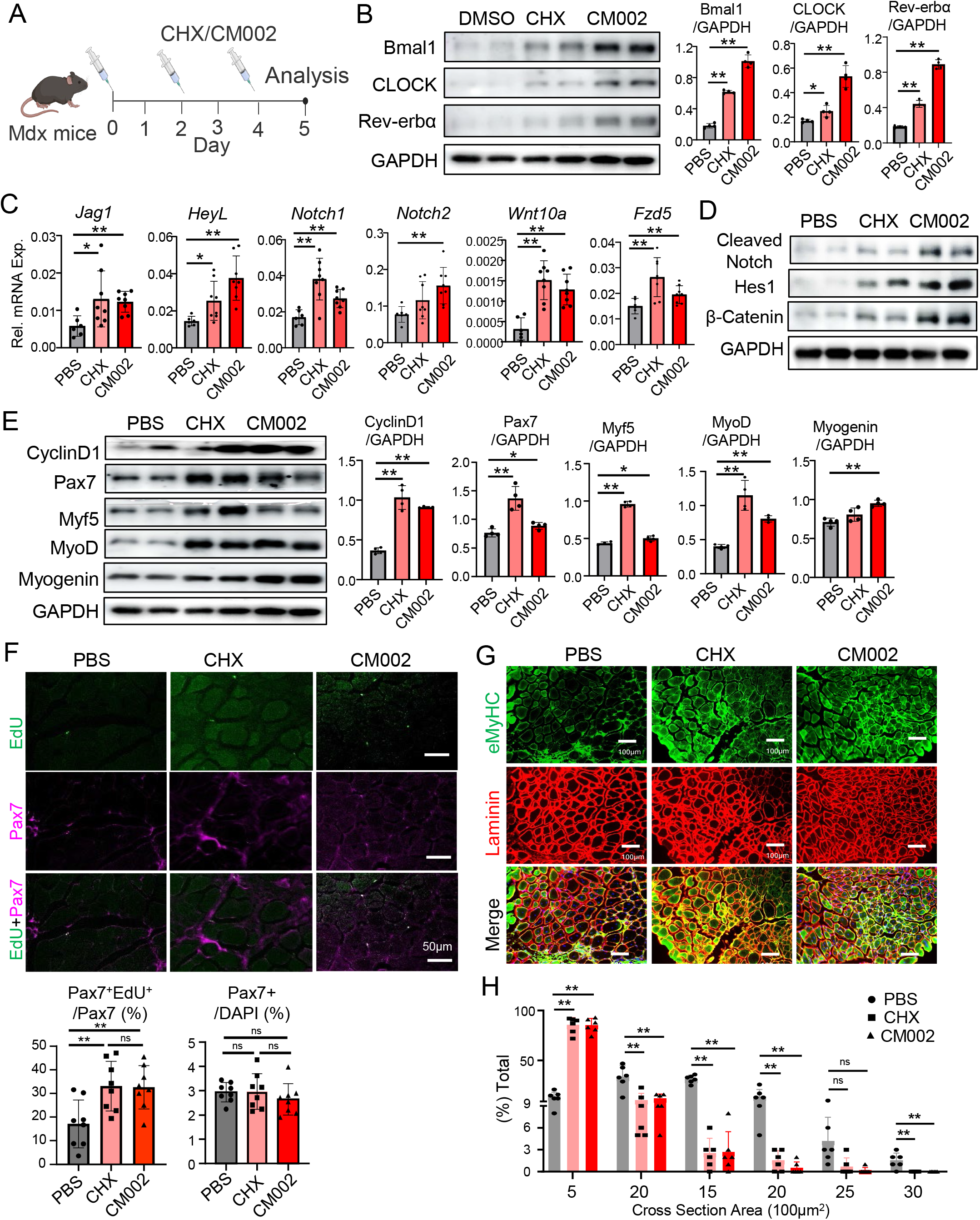
Effect of CM002 and CHX on promoting regenerative myogenesis in dystrophic muscle. (A) Schematic representation of CHX and CM002 treatment time course in mdx mice. Male mdx mice were treated every other day with these compounds (50 μl of 2μM in PBS) for a total of 3 times via intramuscular injection into TA with samples collected after 5 days. (B) Representative images of immunoblot analysis of clock regulators in TA muscle of mdx mice treated with CHX or CM002 with quantitative analysis from three repeats. (C, D) RT-qPCR (C), and immunoblot analysis (D), of Notch and Wnt signaling components in TA muscle of mdx mice with CHX or CM002 treatment. (E) Representative images of immunoblot analysis of regenerative myogenesis in TA of mdx mice treated with CHX or CM002. Each lane represents pooled samples of 3-4mice/lane. (F) Representative images of immunofluorescence Pax7 staining (Magenta) with EdU labeling (green) to determine proliferation in TA of mdx mice. N=8/group. (G, H) Representative images of immunofluorescence staining of embryonic MyHC (green) and laminin (red) of mdx mice TA cross section following CHX or CM002 treatments (G) with quantitative analysis (H). *p<0.05 and ** p<0.01 CM002 or CHX treatment vs. DMSO control by Student’s *t* test.

## Discussion

The immense regenerative potential of muscle stem cells could be exploited for muscle disease applications (Kuang & Rudnicki, 2008, Seale, Polesskaya et al., 2003). Our current study identified a circadian clock-dependent regulatory mechanism that directly controls key components of the Notch and Wnt signaling pathways, critical drivers of muscle regeneration. Our findings demonstrate that the circadian clock orchestrates the temporal activation of these signaling programs to couple progenitor expansion with subsequent myogenic progression during regenerative myogenesis. Mechanistically, clock activation promotes early Notch signaling to support proliferative expansion of myogenic progenitors, while facilitating Wnt signaling at later stages to drive myogenic differentiation. Importantly, reinforcing this endogenous timing mechanism through pharmacological clock activation enhanced MuSC regenerative function in both acute injury and dystrophic muscle, establishing clock-controlled signaling outputs as targetable mechanism for regenerative repair.

Most studies of circadian clock function in skeletal muscle stem cells to date are limited to loss-of-function models. Using a unique model with Bmal1 over-expression selectively in satellite cells, the current study demonstrated the potential for augmenting clock activity to enhance the myogenic properties of MuSCs and its progenitor populations. Notably, MuSC clock gain-of-function did not induce overt phenotypes in skeletal muscle under basal conditions. Upon injury, the peak of regeneration in sBMKI mice was delayed ∼2 days, while a later augmented peak was detected at dpi 5 coupled with an enhanced yet extended regenerative myogenic time course that persisted up to 30 days. This regenerative dynamic induced by clock gain-of-function in MuSCs is accompanied with an early induction of the Notch signaling and enhanced Wnt pathway activity at later time points of regeneration. We have previously identified that signaling components of the Wnt cascade were direct clock targets (Chatterjee et al., 2013, Guo et al., 2012), while new findings revealed that the Notch pathway is also under clock transcriptional control, with Bmal1 chromatin occupancy detected at E-box sites of identified gene promoters, For normal muscle regeneration, a temporal transition from early activation of Notch to later stages of Wnt signaling is required to drive myogenic progression (Brack et al., 2008, Conboy & Rando, 2002, Jin, Kim et al., 2009, Seale et al., 2003). By activating the clock to re-enforce this intrinsic temporal transition from Notch to Wnt, either genetically or pharmacologically, it facilitated a normal temporal course required for myogenic progression to allow proliferative expansion of the MuSC pool prior to myogenic development. The Notch pathway is known to play a crucial role in driving the self-renewal to maintain the satellite cell pool (Conboy & Rando, 2002, Wen, Bi et al., 2012) with its activation displaying context- and developmental stage-dependent effects on myogenesis (Bi, Yue et al., 2016, Peng, Scinto et al., 2025). The increased Notch activation in MuSCs with ectopic clock expression in the early stage of regeneration may have extended proliferative expansion phase with maintenance of self-renewal. Indeed, inhibition of Notch signaling in these cells indeed blocked the increase in their proliferation.

Given the oscillatory nature of molecular clock action via a negative feed-back loop, Bmal1 promotion of Notch signaling and consequent MuSC modulations are likely not persistent, which may have accounted for the disparate effects as compared with constitutive Notch activation in satellite cells (Bi et al., 2016, Wen et al., 2012). Activation of Notch is known to block myogenic differentiation by inhibiting the expression of myogenic factors (Buas, Kabak et al., 2010, Kopan, Nye et al., 1994, Kuroda, Tani et al., 1999). Satellite cell-specific overexpression of the Notch Intracellular Domain, the mediator of Notch signaling following its protease cleavage, blocked myogenic development leading to loss of muscle mass (Wen et al., 2012). The intriguing finding of a delayed but prolonged regenerative time course in the sBMKI could be a result of enhanced Notch activation at early stages, while the subsequent activation of the Wnt pathway that was similarly induced by clock gain-of-function was sufficient to overcome Notch activity to allow the myogenic transition. As a result, a consequently positive impact was evident on regeneration with a moderate myocyte shift toward larger sizes myofibers. It is thus possible that due to the dual transcriptional control of circadian clock on Notch as well as the Wnt pathway, or potentially additional pathways, the impact of clock gain-of-function on regenerative myogenesis is distinct from that of the detrimental effects of promoting Notch or Wnt activity alone (Brack et al., 2008, Brack et al., 2007).

Notch signaling activation is dominant during the early phase of muscle injury and muscle stem cell maintenance. This key pathway in mediating neighboring cell-cell interactions is known to confer cell-autonomous cis-inhibitory effect on signaling within the same cell, while activating intracellular cascade of the receiving cell via trans-activation (Kuintzle, Santat et al., 2025, Sprinzak, Lakhanpal et al., 2010, Thambyrajah, Maqueda et al., 2024). In myogenesis, a recently identified mechanism of MyoD activation of the Notch ligand, Dll1, is involved in inhibiting Notch signaling via a cis-inhibition mechanism (Zhang, Shang et al., 2021). Based on findings of clock target genes involved in the Notch pathway, several mechanisms may account for the self-limited activation observed in sBMKI mice during regeneration. Jag-1, a Notch ligand, was found to be a direct clock target. This ligand is known to induce cis-inhibition in the signal-originating cell itself while stimulating neighboring receiving cells via transactivation (Kuintzle et al., 2025, Thambyrajah et al., 2024). Its regulation in sBMKI cells may elicit divergent signaling outcomes that enable inhibition of Notch signaling propagation during regeneration. A cis-inhibition mechanism mediated by Jag1-Notch 1 in sBMKI-expressing committed myogenic progenitors may lead to the attenuation of Notch signaling with myogenic progression, while allowing nearby MuSC to maintain a self-renewal pool through trans-activation. It is also possible that, as HeyL was also identified as a target of clock regulation, this repressive Notch transcriptional effector, as well as mediator of a negative feedback loop, may contribute to the prolonged but eventual dampening of Notch activation. Consequently, the enhanced myogenic drive induced by clock activity in sBMKI mic was self-limited at the end of regeneration. By re-enforcing an MuSC-intrinsic temporal regulatory element to promote development signaling programs required for normal regenerative myogenesis, clock activation augmented the accumulative myogenic response via coordination of an orderly progression through distinct phases of myogenic activation, stem cell expansion, myogenic commitment and differentiation. Besides Notch and Wnt regulations, circadian clock controls additional developmental signals to fine-tune the homeostatic functions of various stem cell compartments, including the Tgf-β and BMP pathways (Janich et al., 2011, Nam, Guo et al., 2015). It thus remains to be determined whether other clock-controlled pathways could be involved in modulating MuSC behaviors, which may constitute a network of temporally controlled signaling events required for muscle regeneration.

Our current study elucidated clock coordination of both Notch and Wnt activities that facilitated normal myogenesis. During regenerative myogenesis in sBMKI mice, the collective effects of Notch and Wnt signaling cascade driven by clock gain-of-function may underly the prolonged proliferative phase coupled with a delayed transition for nascent myofiber formation. Notch signaling is involved in satellite cell activation and proliferation (Conboy, Conboy et al., 2003, Conboy & Rando, 2002). For a direct test that enhanced Notch activity early during regeneration accounts for the proliferative expansion of sBMKI satellite cells, we obtained primary myoblasts and found that they indeed displayed a higher proliferative rate that was efficiently blocked by inhibition of Notch signaling. Thus, clock-controlled Notch activation underlies the proliferative phenotype in sBMKI cells. This result mirrors our prior finding that impaired Wnt signaling accounts for the defect in myogenic differentiation of Bmal1-null myoblasts (Chatterjee et al., 2013). Additionally, clock activation by CM002 increased Bmal1 chromatin occupancy on Notch pathway gene promoters, providing direct support of Notch pathway gene activation by clock.

Discovery of clock-activating molecules enabled our study of pharmacological clock interventions in muscle regeneration under normal or dystrophic conditions. Chlorhexidine was previously identified as a circadian clock activator with pro-myogenic activities (Kiperman et al., 2023a). We explored its potential application in muscular dystrophy using mdx primary myoblasts as a cellular model, with further demonstration of its clock-dependent actions. In line with Notch and Wnt pathways under the transcriptional control of clock, Bmal1 overexpression or clock activation by CHX or CM002 were able to induce Notch and Wnt genes in myogenic precursors. Importantly, clock regulation of these pathways was applicable to dystrophic disease, as both CHX and CM002 were sufficient to induce respective signaling components in mdx mice. Thus, clock activation of Notch and Wnt pathways may mediate, to a significant extent, the efficacy of clock activators on enhancing regenerative myogenesis, suggesting the potential for pharmacological targeting of this mechanism to promote regeneration. Wnt and Notch abnormalities found in dystrophic muscles contribute to impaired regenerative capacity, a key pathogenic mechanism of loss of muscle mass and function (Den Hartog & Asakura, 2022, Gurriaran-Rodriguez, Kodippili et al., 2024, Tanaka, Kim et al., 2026). Findings of the effect of CHX and CM002 on enhancing regeneration in the acute injury model provided the initial proof-of-principle evidence of targeting the clock for regenerative repair. Importantly, the pro-myogenic effects of clock activators apply to dystrophic disease, with demonstrated efficacy on inducing MuSC proliferation and new myofiber regeneration in the mdx model. Findings of clock function in promoting regenerative myogenesis established a basis for disease interventions, and the ability of CHX and CM002 to induce regenerative repair in the dystrophic condition supports their potential for future therapeutic applications.

Collectively, our studies uncovered a circadian clock-controlled mechanism that coordinates the temporal transition from Notch to Wnt signaling during regenerative myogenesis. These findings define the circadian clock as an upstream transcriptional regulator of developmental signaling networks that govern MuSC regenerative capacity that is therapeutically applicable to dystrophic disease. While significant challenges remain for clinical application of the identified clock-activating compounds, including the development of drug-like leads and evaluation of systemic delivery strategies to achieve functional outcomes, our work established a mechanistic foundation for targeting circadian clock to promote regenerative repair in muscular dystrophy.

## Acknowledgements

KM is a faculty member supported by the NCI-designated Comprehensive Cancer Center at the City of Hope National Cancer Center. This project was supported by National Institute of Health grants R01DK137515 and R56AG080294, and an Innovative Pilot Award from Arthur Riggs Diabetes and Metabolism Research Institute to KM. TK is funded by a predoctoral fellowship from the California Institute for Regenerative Medicine. VY is supported by National Institute of Health grants R01DK128972 and R01DK130499. These funders had no role in study design, data collection and analysis, decision to publish, or preparation of the manuscript.

## CRediT Author contributions statement

XX, JP, TK: data curation and investigation, methodology, formal analysis, manuscript editing; DH and VY: Software, data curation, manuscript review and editing; KM: formal analysis, project administration, manuscript writing and editing, and funding acquisition.

## Ethics approval and consent to participate

All animal experiments in this study were approved by the Institutional Animal Care & Use Committee (IACUC) of City of Hope according to the approval. The protocol number: 17110, entitled “circadian regulation of metabolism” with approval date is from 12/11/2023 to 12/10/2026.

## Disclosure and Competing Interests Statement

The authors declare that they have no conflict of interest that exists related to the subject matter or materials included in this work.

## Data Availability

All data generated and analyzed during this study are included in this published article and associated Supplementary Information files. Additional data that supports the findings of this study are available from the corresponding author upon request.

## Declaration of Interest

None

## Notes

### Competing Interest Statement

The authors have declared no competing interest.

